# Cholesterol Metabolism in a Murine Model of Vertical Sleeve Gastrectomy

**DOI:** 10.1101/2025.07.03.659993

**Authors:** Emily Whang, Alessandra Ferrari, John Paul Kennelly, Yajing Gao, Xu Xiao, Kelsey Jarrett, Alexander Bedard, Thomas Q de Aguiar Vallim, Peter Tontonoz

## Abstract

The most effective and durable treatment for obesity is metabolic bariatric surgery. Currently, vertical sleeve gastrectomy (VSG) has become the preferred surgical method to achieve sustained weight loss and reverse obesity-related disease, including dyslipidemias. Human studies indicate that 40% of VSG patients can discontinue statin treatment by 6 months, while alterations in plasma bile acids and synthesis have been observed after bariatric surgery. Although it is well known that cholesterol is the precursor for all bile acids, the link between changes in plasma cholesterol and bile acids remains unclear in the context of VSG. Here, we present a murine model of VSG that leads to a marked decrease in plasma cholesterol levels despite no difference in body weight compared to sham surgery. Our results demonstrate that gastric resection lowers systemic cholesterol burden by promoting the hepatobiliary elimination of cholesterol through the upregulation of bile acid synthesis. These observations imply that gastric resection impacts overall hepatic physiology and cholesterol homeostasis independent of weight loss.

## Introduction

Children are becoming overweight at younger ages, and childhood obesity, along with its co-morbidities, is rising at an alarming rate.^1–2^ Since the 1980s, the rate of childhood obesity has tripled in the US, while the prevalence in adolescents has quadrupled.^3–4^ Several studies have now linked excess childhood weight to the development of type 2 diabetes mellitus, dyslipidemias, cardiovascular disease, and metabolic dysfunction associated steatotic liver disease (MASLD) in adulthood.^5–9^ Currently, the most effective and durable treatment for obesity is metabolic bariatric surgery, which includes Roux-en-Y gastric bypass and vertical sleeve gastrectomy (VSG). RYGB is a restrictive and malabsorptive procedure in which a small gastric pouch is created with diversion of food into the distal jejunum, while VSG removes 70-80% of the stomach. Based on its simplicity and low complication rate, VSG has now become the preferred surgical method to achieve sustained weight loss and reverse obesity-related diseases.

The efficacy of VSG in humans is well-established, but the mechanisms driving its effects are not fully understood. This is due, in part, to the multifaceted physiologic changes that involve various organs and metabolic pathways after VSG. Many groups have focused on understanding how VSG contributes to weight loss and the resolution of diabetes, but few have studied the impact of VSG on dyslipidemias despite human studies indicating that 40% of VSG patients can discontinue statin treatment by 6 months.^10–11^ Furthermore, alterations in plasma bile acids and synthesis have been observed after bariatric surgery.^12–14^ Although it is well known that cholesterol is the precursor for all bile acids, the link between changes in plasma cholesterol and bile acids remains unclear in the context of VSG.

While previously thought to be a restrictive process, recent studies indicate that the beneficial effects of VSG do not exclusively depend on weight loss and caloric restriction.^15–16^ For instance, diabetic patients can stop medication within days of surgery even before weight loss occurs.^17^ In mice, hepatic steatosis resolves after VSG, but not in pair-fed, weight-matched mice.^18^ These observations imply that gastric resection impacts overall hepatic physiology and lipid metabolism independent of weight loss.

Here we present a murine model of VSG that leads to a marked decrease in plasma cholesterol levels despite no difference in body weight compared to sham surgery. VSG mice also exhibit reduced hepatic cholesterol, liver steatosis, and hepatic fibrosis markers independent of weight loss. To elucidate the mechanism of VSG’s cholesterol-lowering effect, we characterized the influx and efflux pathways responsible for cholesterol homeostasis by assessing food intake, cholesterol absorption, hepatic lipoprotein uptake, and hepatobiliary cholesterol excretion. Our findings conclude that gastric resection reduces systemic cholesterol burden by promoting the hepatobiliary elimination of cholesterol through the upregulation of bile acid synthesis.

## Results

### VSG lowers plasma cholesterol independent of weight loss

In our murine model of VSG, 8-week-old male mice were fed a western diet containing 0.2% cholesterol (RD D12079B) for 8 weeks, then matched for overall body weight and growth velocity (average weekly weight gain) before sham or VSG surgery. Sham surgery was performed by opening the peritoneum and clamping the stomach for a few minutes without resection, while VSG was achieved by suturing the gastric mucosa to create a tubular remnant after resecting 70-80% of the stomach. Prior to surgery, all mice underwent an overnight fast. After surgery, both sham and VSG mice received a liquid diet for 3 days, then resumed a western diet containing 0.2% cholesterol. All mice received baytril-treated water for 5 days post-operatively.

VSG mice displayed similar body weights compared to sham mice on a western diet containing 0.2% cholesterol (Fig. 1A). Both groups lost weight initially post-surgery. Sham mice regained their body weights by week 5, while VSG mice recovered their body weights by week 6 (Fig. 1B). Weights were not different between groups thereafter, yet plasma cholesterol at week 6 and 11 were reduced in VSG mice (Fig. 1C-D). The effect on plasma cholesterol was sustained at week 14 on a western diet containing 1.25% cholesterol (Fig. 1E), with decreased low-density lipoprotein (LDL) and high-density lipoprotein (HDL) cholesterol levels in VSG mice (Fig. 1F). This was consistent with decreased serum ApoB100 after VSG (Fig. 1G). No difference was appreciated in serum ApoA1 or ApoE levels (Fig. S1A). Plasma triglycerides were slightly higher in VSG mice 6 weeks post-operatively, but were not different compared to sham at week 11 (Fig. 1H-I).

**Fig. 1.**
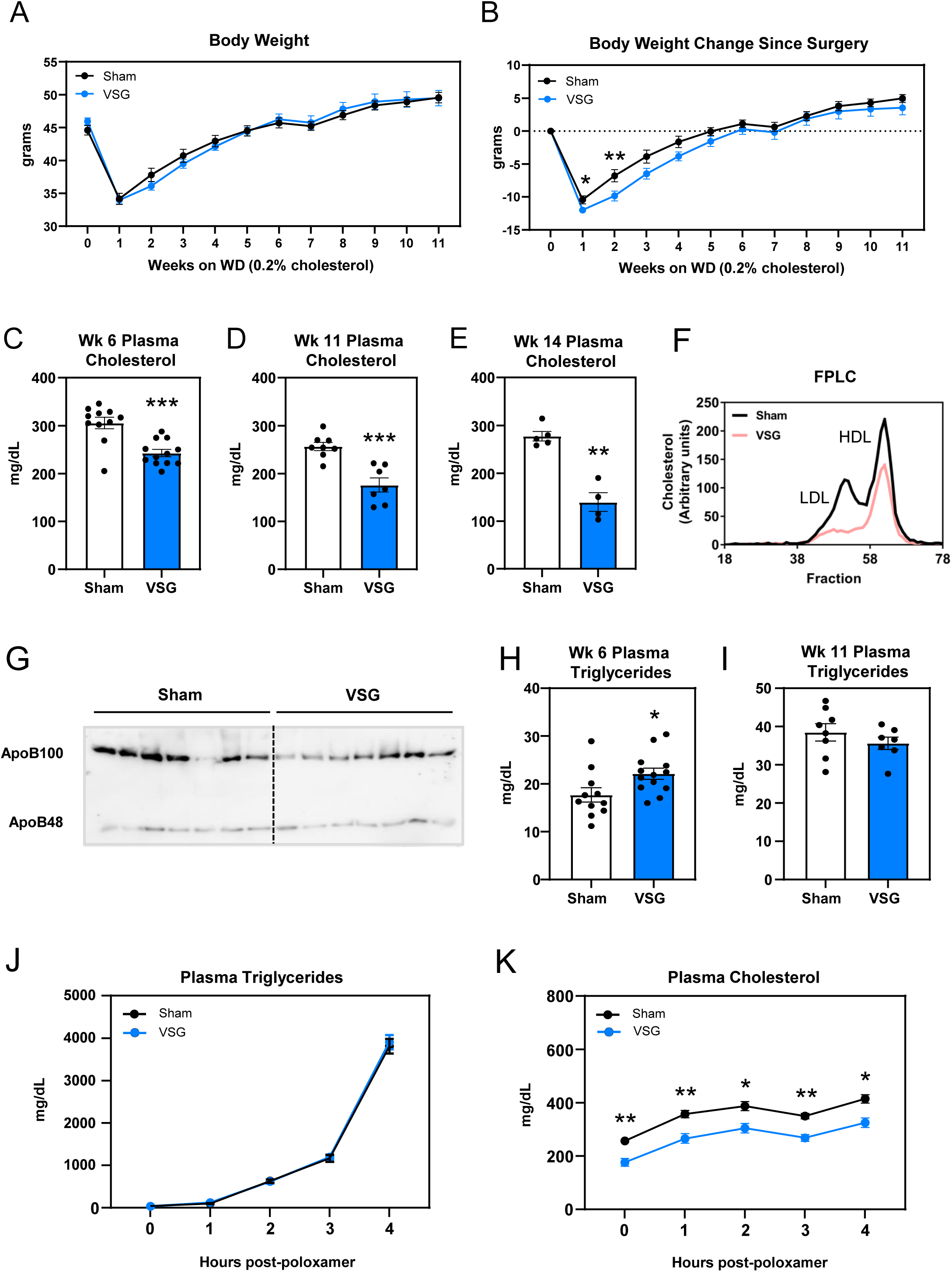
VSG decreases plasma cholesterol independent of weight loss. (A) Post-op gross body weight of sham and VSG mice on western diet containing 0.2% cholesterol (WD), n = 10 vs 12; (B) Body weight change since surgery of sham and VSG mice on WD, n = 10 vs 12; (C) Week 6 non-fasting plasma cholesterol in sham and VSG mice on WD, n = 11 vs 12, p = 0.0007; (D) Week 11 fasting plasma cholesterol in sham and VSG mice on WD, n = 8 vs 7, p = 0.0012; (E) Week 14 fasting plasma cholesterol in sham and VSG mice on WD + 1.25% cholesterol, n = 5 vs 4, p = 0.0159; (F) Week 14 fasting LDL-cholesterol and HDL-cholesterol in sham and VSG mice on WD 1.25% cholesterol, pooled samples, n = 5 vs 4; (G) Week 6 non-fasting serum ApoB100/48 in sham and VSG mice on WD, n = 7 vs 7; (H) Week 6 non-fasting plasma triglyceride levels in sham and VSG mice on WD, n = 10 vs 12, p = 0.0291; (I) Week 11 fasting plasma triglyceride levels in sham and VSG mice on WD, n = 8 vs 7; (J) Week 11 plasma triglyceride levels in sham and VSG mice after intraperitoneal injection with Poloxamer, n = 8 vs 7; (K) Week 11 plasma cholesterol levels in sham and VSG mice after intraperitoneal injection with Poloxamer, n = 8 vs 7. (*P < 0.05, ** P < 0.01, *** P < 0.001, ****P < 0.0001)

To determine whether altered VLDL secretion could explain the differences in plasma lipoproteins, mice were injected with the lipoprotein lipase inhibitor Poloxamer-407, and blood was collected hourly for up to 4 hours. The rate of VLDL secretion of both triglycerides and cholesterol were similar between groups, though plasma cholesterol was decreased at all time points in VSG mice (Fig. 1J-K).

### VSG does not reduce dietary cholesterol intake

Since systemic cholesterol homeostasis is achieved by balancing cholesterol influx and efflux, we first characterized cholesterol influx pathways, including food intake and absorption, in sham and VSG mice. All experiments were performed in mice on a western diet containing 0.2% cholesterol. Gross food intake was lower in VSG mice between weeks 1 and 2, but not different compared to sham for the remaining weeks (Fig. 2A). The reduction in food intake correlated with a decrease in weight gain during the first 2 weeks in VSG mice (Fig. 2B). Otherwise, weekly weight gain was not different between groups, suggesting intact absorption after VSG. Furthermore, there was no difference in intestinal histology, intestinal length, or fecal weights between sham and VSG mice (Fig. 2C-E). Fatty acid absorption, assessed with a 5% sucrose polybehenate diet, was also not different between sham and VSG mice (Fig. 2F, S1B).

**Fig 2.**
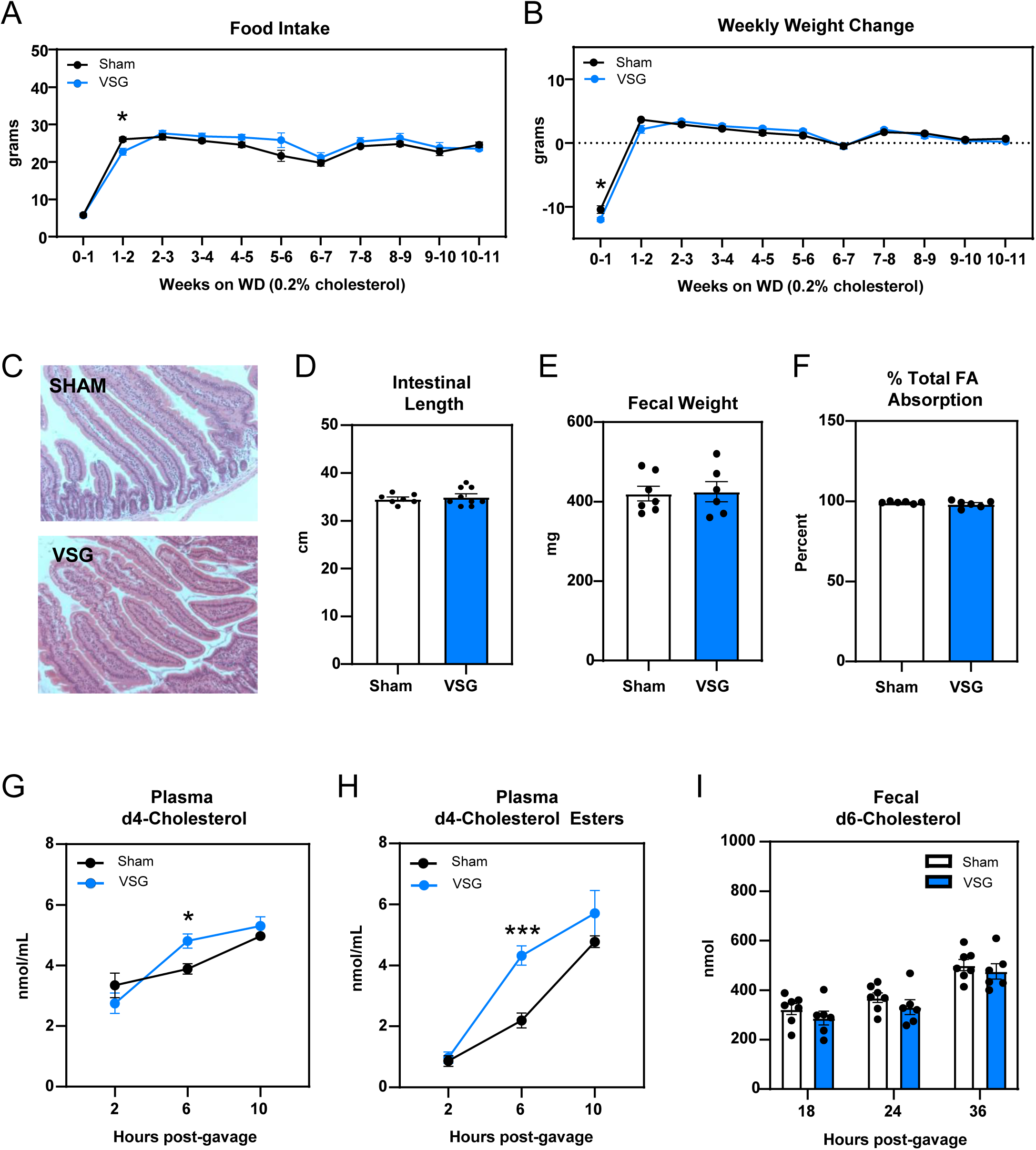
VSG does not reduce dietary cholesterol intake. (A) Food intake of sham and VSG mice on WD, n = 10 vs 12; (B) Weekly weight gain of sham and VSG mice on WD, n = 10 vs 12; (C) Representative H and E staining of jejunal tissue in sham and VSG mice; (D) Intestinal length, n = 7 vs 9, p = 0.9353; (E) Fecal weights, n = 7 vs 6, p = 0.8755; (F) % fatty acid absorption in sham and VSG mice on a 5% sucrose polybehenate diet at 6 weeks post-op, n = 6 vs 5; (G) Plasma levels of deuterated cholesterol after after gavage with deuterium labeled cholesterol-25,26,26,26-d4 cholesterol (D4-cholesterol). Performed at 6 weeks post-op, n = 8 vs 8, p = 0.6251, 0.0235, 0.7313; (H) Plasma levels of deuterated cholesterol esters after gavage with deuterium labeled cholesterol-25,26,26,26-d4 cholesterol (d4-cholesterol). Performed at 6 weeks post-op, n = 8 vs 8, n = 8 vs 8, p = 0.9611, 0.0004, 0.5936; (I) Fecal levels of deuterated cholesterol after gavage with deuterium labeled cholesterol-2,2,3,4,4,6-d6 cholesterol (d6-cholesterol). Performed at 6 weeks post-op. n = 7 vs 6, p = 0.7132, 0.6908, 0.8951. (*P < 0.05, ** P < 0.01, *** P < 0.001, ****P < 0.0001)

To evaluate cholesterol absorption, we measured the appearance of orally administered deuterium labeled cholesterol-25,26,26,26-d4 (d4-cholesterol) in the blood of sham and VSG mice. Plasma levels of d4-cholesterol and d4-cholesterol esters were not decreased in VSG mice compared to sham mice (Fig. 2G-H), consistent with intact cholesterol absorption after VSG. Interestingly, VSG mice displayed an increase in plasma d4-cholesterol and d4-cholesterol esters at 6 hours post-gavage, suggesting more efficient absorption. In a separate experiment, cholesterol absorption was assessed by measuring the loss of deuterated cholesterol-2,2,3,4,4,6-d6 (d6-cholesterol) in feces after oral gavage. Sham and VSG showed no difference in fecal d6-cholesterol loss (Fig. 2I), confirming that decreased cholesterol absorption is not responsible for the reduction in plasma cholesterol levels after VSG.

### VSG improves hepatic steatosis independent of weight loss

The liver plays an integral role in cholesterol homeostasis by balancing exogenous intake, de-novo synthesis, and hepatobiliary excretion. This process is made possible through reverse cholesterol transport, a process in which HDL transports peripheral cholesterol to the liver for elimination. Despite no difference in dietary cholesterol intake, VSG mice revealed lower hepatic cholesterol levels, while liver triglyceride levels were unchanged compared to sham (Fig. 3A-B). Therefore, to assess whether this was due to impaired lipoprotein uptake by the liver, we performed tracer studies using radiolabeled HDL containing [^14^C]-cholesterol. After intravenous injection of [^14^C]-HDL cholesterol into sham and VSG mice, we evaluated hepatic clearance of [^14^C] from the blood over 1 hour (Fig. 3C). There was no difference in the rate of hepatic uptake between groups although plasma [^14^C] levels were marginally lower in VSG mice compared to sham mice in this experiment.

**Fig. 3.**
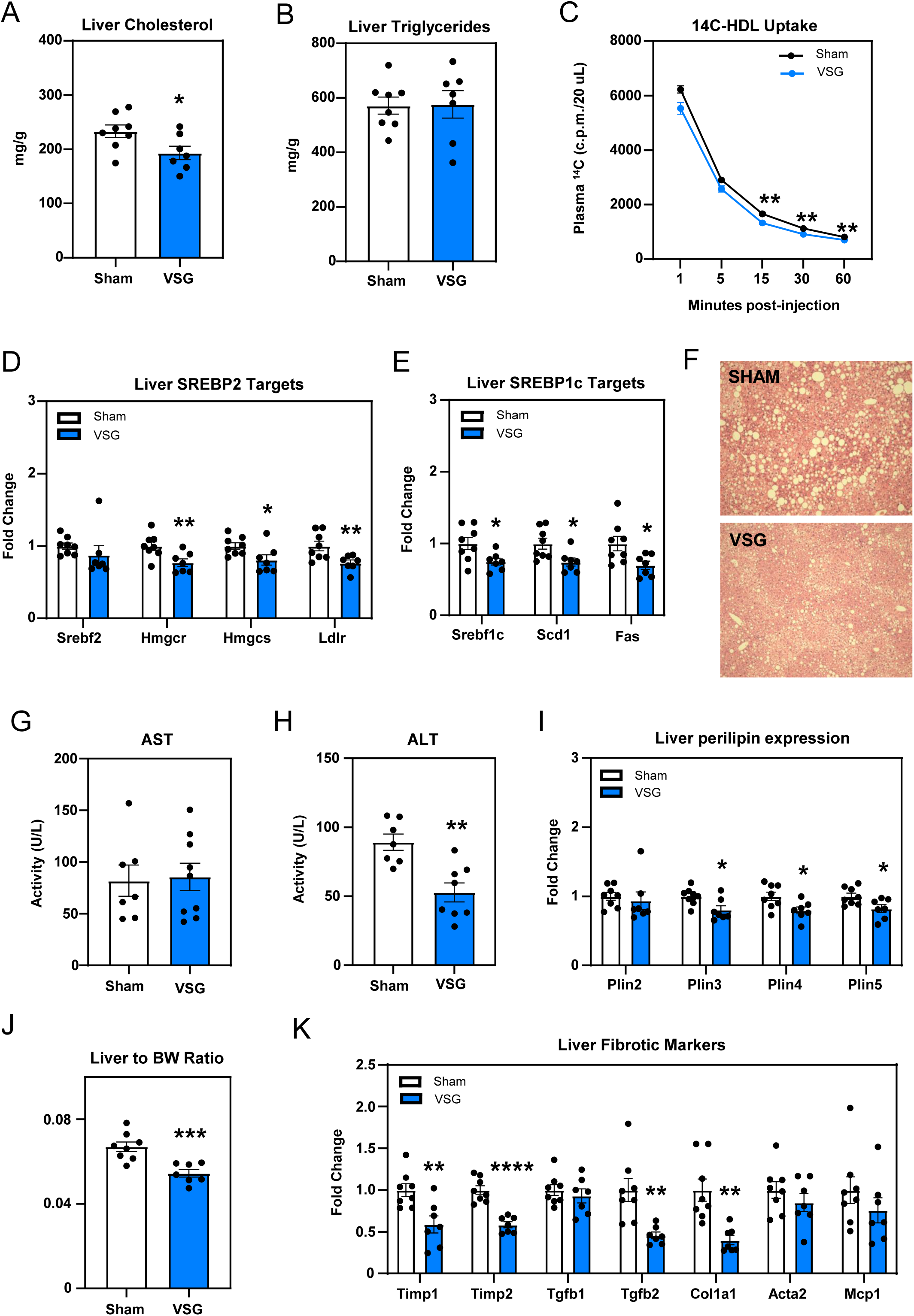
VSG improves hepatic steatosis independent of weight loss. (A) Liver cholesterol 11 weeks on WD post-op sham vs VSG, n = 8 vs 7, p = 0.0365; (B) Liver triglycerides 11 weeks on WD post-op sham vs VSG, n = 8 vs 7, p = 0.9370; (C) Plasma levels of 14C after injection of 14C-HDL in sham vs VSG mice, n = 7 vs 9, p = 0.0761, 0.1341, 0.0043, 0.0011, 0.0054; (D) Gene expression of Srebp2 target genes in the livers of sham vs VSG mice at 7 weeks post-op, n = 8 vs 7, p = 0.390777, 0.011642, 0.046024, 0.011987; (E) Gene expression of Srebp1c target genes in the livers of sham vs VSG mice at 7 weeks post-op, n = 8 vs 7, p = 0.025475, 0.017420, 0.024693; (F) Representative H and E staining of jejunal tissue in sham and VSG mice at 14 weeks post-op; (G) Serum AST at 7 weeks post-op, n = 7 vs 8, p = 0.8546; (H) Serum ALT at 7 weeks post-op, n = 7 vs 8, p = 0.0015; (I) Gene expression of hepatic perilipins in sham vs VSG mice, n= 8 vs 7; (J) Liver to body weight ratio at 11 weeks post-op, n = 8 vs 7, p = 0.0009; (I) Gene expression of hepatic fibrotic markers at 11 weeks post-op, n = 8 vs 7, p = 0.007649, 0.000022, 0.520544, 0.004862, 0.002223, 0.322249, 0.289366. (*P < 0.05, ** P < 0.01, *** P < 0.001, ****P < 0.0001)

Upon delivery to the liver, cholesterol may be utilized, stored, distributed to peripheral organs, or eliminated in the biliary system. We have previously shown that a substantial fraction of lipoprotein-derived cholesterol in the liver is transported to the endoplasmic reticulum (ER) before excretion into the hepatobiliary system.^19^ When ER cholesterol is low, de novo cholesterol synthesis is increased through transcriptional up-regulation of the SREBP2 pathway. Therefore, changes in the expression of SREBP2 targets provide a reflection of ER cholesterol content. Although overall hepatic cholesterol was decreased after VSG, genes involved in de-novo cholesterol synthesis were suppressed in the livers of VSG mice compared to sham mice (Fig. 3D), suggesting an increase in cholesterol trafficking to the ER. Hepatic expression of SREBP1c targets was also down-regulated after VSG, signifying a decrease in fatty acid synthesis (Fig. 3E).

Although liver triglyceride content was not different between groups, VSG mice demonstrated reduced hepatic steatosis after 14 weeks on a high fat/high cholesterol diet despite having similar weights to sham mice (Fig. 3F). This finding correlated with decreased serum transaminases (Fig. 3G-H), lower liver to body weight ratios (Fig. 3J), and a decrease in the expression of hepatic fibrotic markers(Fig. 3K) after VSG compared to sham surgery. VSG mice also demonstrated a decrease in hepatic perilipin expression compared to sham mice (Fig. 3I), suggesting decreased lipid droplet formation. These findings, along with lower hepatic cholesterol levels, suggest that VSG induces a metabolic shift toward cholesterol efflux in the liver.

### VSG promotes hepatobiliary cholesterol elimination by increasing conversion to bile acids

Having shown above that cholesterol influx was not different between sham and VSG mice, we hypothesized that plasma and hepatic cholesterol levels could be lower in VSG as a result of increased hepatobiliary elimination of cholesterol. This could occur through secretion of cholesterol itself by the ABCG5/8 transporter and/or by conversion to bile acids via CYP7A1. Consistent with our hypothesis of cholesterol efflux, hepatic expression of both ABCG5/8 and CYP7A1 were higher in VSG mice versus sham mice (Fig. 4A). Gallbladder volumes were also elevated in VSG mice (Fig. 4C). Although biliary cholesterol concentration was not increased, total biliary cholesterol was increased due to an increase in gallbladder volume after VSG (Fig. 4D-E). However, total cholesterol levels in the feces were lower in VSG mice (Fig. 4F), demonstrating that increased hepatobiliary excretion of free cholesterol was not responsible for low plasma and hepatic cholesterol levels after VSG. These results suggested that the increase in cholesterol efflux was due to an increase in conversion to bile acids.

**Fig. 4.**
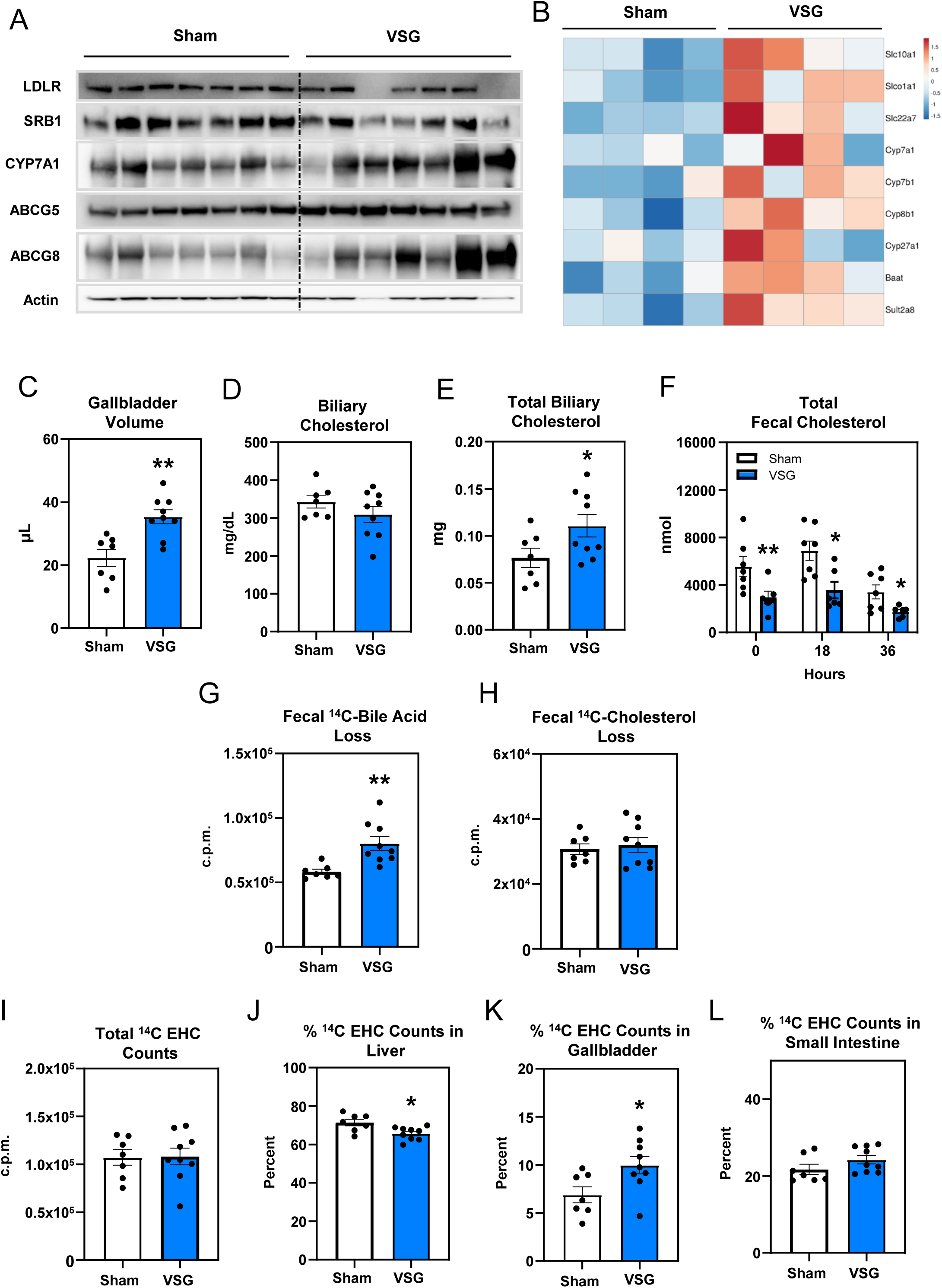
VSG promotes the conversion of cholesterol to bile acids. (A) Immunoblot of liver LDLR, SRB1, CYP7A1, ABCG5/8 expression at 11 weeks post-op; (B) Heatmap showing expression of genes involved in bile acid transport and synthesis in the liver at 11 weeks post-op. Slc10a1, Slco1a1, Slc22a7, Cyp8b1, Baat, and Sult2a8 achieved significance, n = 8 vs 7; (C) Gallbladder volume at 7 weeks post-op, n = 7 vs 9, p = 0.0024; (D) Biliary cholesterol concentration at 7 weeks post-op, n = 7 vs 9, p = 0.2323; (E) Total cholesterol content in GB at 7 weeks post-op, n = 7 vs 9, p = 0.0478; (F) Fecal cholesterol content over 36 hours at 6 weeks post-op, n = 7 vs 6, p = 0.01399, 0.02215, 0.03497; (G) Fecal loss of 14C-cholesterol as bile acids over 3 days after intravenous injection of [14C]-labeled HDL at 7 weeks post-op, n = 7 vs 9, p = 0.0032; (H) Fecal loss of 14C-cholesterol as free cholesterol over 3 days after intravenous injection of [14C]-labeled HDL, n = 7 vs 9, p = 0.6331; (I) Total [14C] counts within the liver, GB, and small intestine 3 days after intravenous injection of [14C] labeled HDL, n = 7 vs 9, p = 0.9314; (J) % counts in the liver, n = 7 vs 9, p = 0.0201; (K) % counts in the GB, n = 7 vs 9, p = 0.0239; (L) % counts in the intestine, n = 7 vs 9, p = 0.1654. (*P < 0.05, ** P < 0.01, *** P < 0.001, ****P < 0.0001)

In line with this model, hepatic RNA-sequencing analysis demonstrated changes in bile acid metabolism, including an increase in genes involved in bile acid synthesis in VSG versus sham mice (Fig. 4B). To show that VSG promotes the conversion of cholesterol to bile acids, we performed radiolabeled tracer studies using HDL containing [^14^C]-cholesterol to characterize hepatobiliary cholesterol loss in the feces. After intravenous injection of [^14^C]-HDL cholesterol into sham and VSG mice, we measured [^14^C] excretion into the feces over 3 days and fractionated the aqueous (bile acids) phase from the lipid (cholesterol) phase in the feces. We observed an increase in [^14^C] within the bile acid fecal compartment of VSG mice versus sham mice, but no difference in the fecal lipid (cholesterol) fraction over 3 days (Fig. 4G-H). This confirmed that cholesterol was being converted to bile acids at an increased rate after VSG.

3 days after injection with HDL containing [^14^C]-cholesterol, mice were sacrificed and the liver, gallbladder, and small intestine were collected to measure [^14^C] counts (Fig. S2). Total radioactivity was measured within the enterohepatic circulation (EHC) and expressed as the sum of counts in the entire liver, gallbladder, and small intestine. Total [^14^C] EHC counts were not different between VSG and sham mice (Fig. 4I). However, the percentage of [^14^C] counts in the EHC was decreased in the liver and increased in the gallbladder of VSG mice (Fig. 4J-K), which is consistent with an increase in the hepatobiliary elimination of [^14^C]-cholesterol. There was no difference in the percentage of [^14^C] counts in the small intestine (Fig. 4L).

### VSG promotes bile acid flux out of the liver

Despite an increase in bile acid synthesis, we found that intrahepatic bile acids were low after VSG, particularly among conjugated bile acids (Fig. 5A-C). This was characterized by reductions in taurocholic acid (TCA) and tauro-β-muricholic acid (Fig. S3-4). To determine whether this reflected bile acid loss from impaired bile acid absorption, we measured the appearance of orally administered [^14^C]-TCA in the blood of sham and VSG mice. Although VSG mice demonstrated a much more varied curve, the absorption of [^14^C]-TCA was not decreased after VSG compared to sham (Fig. 5D). In fact, [^14^C] counts appeared to be elevated at all time points in VSG mice, though the area under the curve was not statistically significant (Fig. 5E). There was also no difference in the loss of [^14^C]-TCA in the feces collected over 24 hours (Fig. 5F) and 72 hours (Fig. S5A) post-gavage between sham and VSG mice. These findings indicate that the decrease in intrahepatic bile acids after VSG is not due to impaired TCA absorption.

**Fig. 5.**
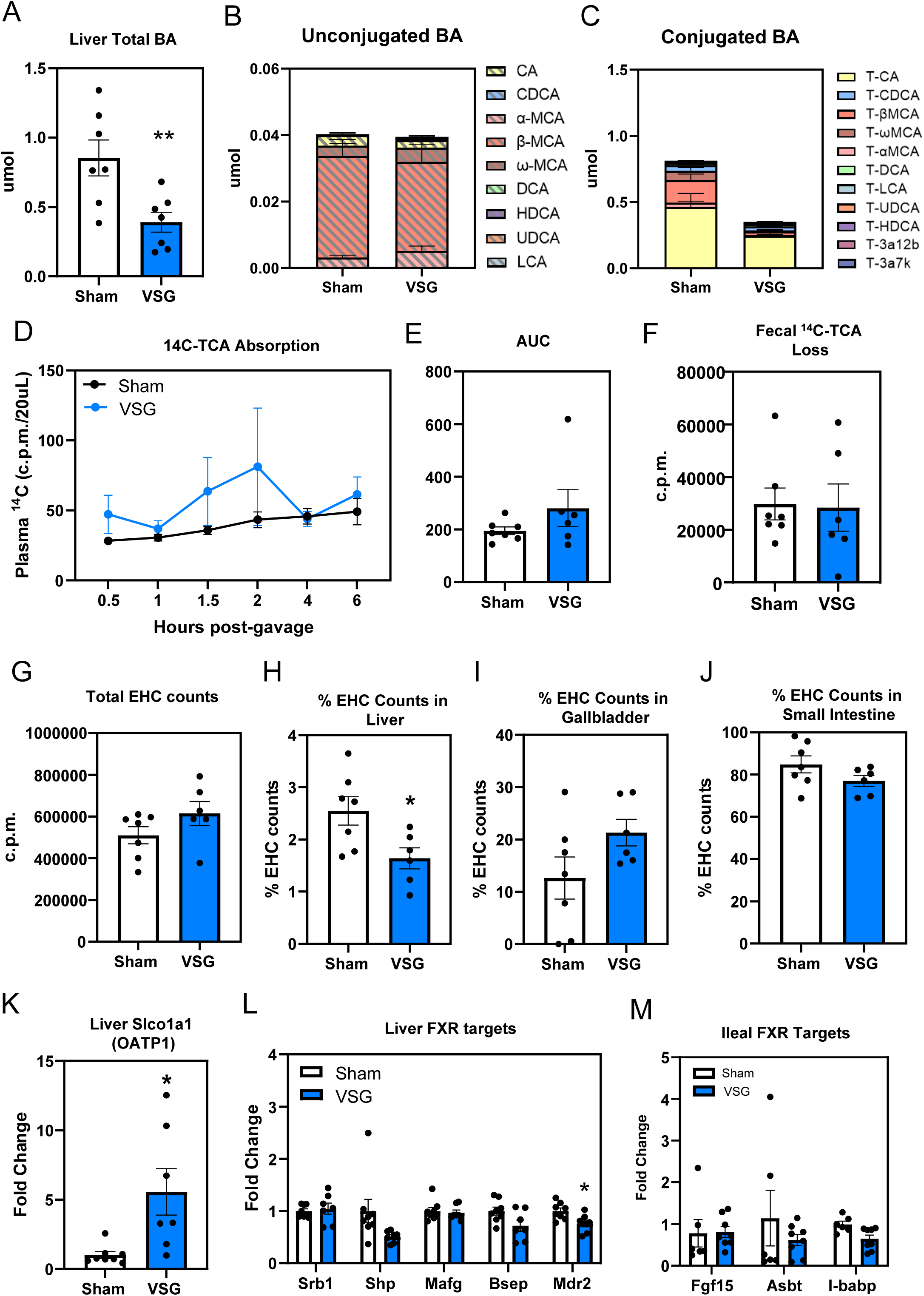
VSG promotes bile acid flux out of the liver. (A) Total liver bile acids at 11 weeks post-op, n = 7 vs 7, p = 0.0088; (B) Unconjugated liver bile acids at 11 weeks post-op, n = 7 vs 7; (C) Conjugated liver BA at 11 weeks post-op, n = 7 vs 7; (D) Plasma levels of 14C after gavage with 14C-taurocholic acid, n = 7 vs 6; (E) AUC of plasma levels of 14C after gavage with 14C-taurocholic acid, n = 7 vs 6, p = 0.2833; (F) Fecal14C counts over 24 hours after gavage with14C-taurocholic acid, n = 7 vs 6, p = 0.7861; (G) Total 14C counts in the liver, GB, and small intestine after gavage with 14C-taurocholic acid, n = 7 vs 6, p = 0.1717; (H) % counts in the liver (7-2), n = 7 vs 6, p = 0.0219; (I) % counts in the GB (7-2), n = 7 vs 6, p = 0.0975; (J) % counts in the intestine (7-2), n = 7 vs 6, p = 0.1339; (K) Hepatic gene expression of Slco1a1 at 11 weeks post-op, n = 8 vs 7, p = 0.0344; (L) Hepatic FXR gene targets at 11 weeks post-op, n = 8 vs 7, Srb1 (p = 0.69455), Shp (p = 0.06675), Mafg (p = 0.7259), Bsep (p = 0.05594), Mdr2 (p = 0.01816); (M) Ileal FXR gene targets at 7 weeks post-op, n = 6 vs 9. (*P < 0.05, ** P < 0.01, *** P < 0.001, ****P < 0.0001)

3 days after [^14^C]-TCA gavage, we measured total [^14^C] counts within the enterohepatic circulation (EHC) as the sum of counts in the liver, gallbladder, and intestine (Fig. S5B-D). The percentage of [^14^C] counts in each tissue compartment relative to the total EHC [^14^C] counts was calculated for each mouse. Due to their hepatotoxic effects, bile acids are typically present in very low quantities within the liver. Although total EHC [^14^C] counts were not different in VSG and sham mice, the percentage of [^14^C]-TCA in the liver was lower in VSG mice, consistent with the observation that VSG mice have reduced intrahepatic bile acids (Fig. 5G-H). There was also a shift towards an increase in the percentage of [^14^C]-TCA within the gallbladder, consistent with increased bile acid export into the biliary system (Fig. 5I). Total [^14^C]-TCA counts in the gallbladder, calculated as counts per microliter multiplied by gallbladder volume, was also noted to be higher in VSG mice. There was no difference in the percentage of [^14^C]-TCA in the small intestine (Fig. 5J). These findings suggest an increase in bile acid flux out of the liver and into the biliary system.

We considered that the decrease in intrahepatic bile acids after VSG could be a result of decreased hepatic bile acid uptake. However, the expression of *Slco1a1* (OATP1) and *Slc10a1* (NTCP) were upregulated in the livers of VSG mice (Fig. 5K, S6), suggesting increased hepatic bile acid uptake. Decreased intrahepatic bile acids in the setting of increased uptake and bile acid synthesis points to an overall shift towards increased bile acid efflux and enterohepatic cycling after VSG, consistent with the results from our radiolabeled tracer assay using [^14^C]-taurocholic acid.

Bile acids circulate from the liver through the biliary system into the small intestine, where they facilitate lipid absorption. In the terminal ileum, 95% of bile acids are reabsorbed and transported back to the liver for recycling. The subsequent return of bile acids to the liver suppresses bile acid synthesis to prevent overproduction via hepatic and intestinal farnesoid X receptor (FXR). We found that the reduction in intrahepatic bile acids after VSG correlated with a trend towards decreased expression of hepatic FXR targets in VSG mice (Fig. 5L). This is consistent with an increase in bile acid synthesis secondary to decreased FXR activity and loss of negative feedback inhibition by *Shp.* However, *Cyp7a1* was not consistently elevated on a transcriptional level after VSG (Fig. S6). Therefore, it is unclear whether alterations in the FXR pathway contribute to overall bile acid synthesis or if the decrease in FXR activity merely reflects lower intrahepatic bile acids after VSG.

Upon bile acid absorption, the activation of intestinal FXR upregulates *Fgf15* expression to suppress bile acid synthesis and down-regulates *Asbt* expression to prevent further bile acid absorption. Thus, decreased bile acid absorption manifests as a decrease in *Fgf15* expression and increase in *Asbt* expression in the ileum. After VSG, there was no difference in the expression of *Fgf15* or *Asbt* (Fig. 5M), which is consistent with our results demonstrating that VSG does not impair bile acid absorption. These findings also suggest that the increase in bile acid synthesis in VSG mice is not regulated by changes in intestinal FXR activity and FGF15 signaling.

## Discussion

Here we present a murine model of VSG to demonstrate the weight independent effects of gastric resection on overall cholesterol homeostasis and bile acid metabolism. Our results indicate that VSG promotes the hepatobiliary elimination of cholesterol by increasing the conversion of cholesterol to bile acids, thereby decreasing plasma and hepatic cholesterol levels independent of weight loss. This is consistent with other murine models of VSG, where plasma cholesterol levels are reduced, while serum bile acids increase.^20–22^ Although these findings are reminiscent of Cyp7a1 transgenic mice ^23^, evidence for increased bile acid synthesis following VSG has been overshadowed by the variability in *Cyp7a1* expression in prior studies. Most groups have relied on gene expression alone without evaluating protein expression. However, in our study, we noted that CYP7A1 protein levels were consistently elevated in VSG mice compared to sham, while *Cyp7a1* mRNA levels were highly variable. Thus, protein quantification of CYP7A1 may serve as a more reliable indicator of bile acid synthesis than gene expression, though the cause for this discrepancy remains unclear.

To better understand the VSG-induced physiologic changes that increase cholesterol conversion to bile acids, we investigated known drivers of bile acid synthesis, such as loss of negative feedback inhibition from bile acid loss and/or decreased FXR signaling. Bile acid loss due to malabsorption was not observed in VSG mice based on radiolabeled tracer studies with [^14^C]-taurocholic acid. Our study, however, was limited to taurocholic acid alone. Thus, we are unable to comment on the absorption of other bile acid species, though the ileal expression of *Fgf15* and *Asbt* was unchanged after VSG, which is consistent with intact bile acid absorption. On the other hand, it is possible that gastric resection may increase bile acid loss due to increased enterohepatic cycling as 5% of bile acids are lost with each cycle. However, VSG mice did not display an overall decrease in [^14^C] counts within the liver, gallbladder, and intestine after gavage with [^14^C]-taurocholic acid. This is inconsistent with increased loss of bile acids, though we are unable to exclude the possibility of miniscule losses that may only be detected over time. Therefore, further assays will need to be performed to determine whether VSG induces bile acid loss.

Our findings do not support the idea that VSG-induced bile acid synthesis is related to suppression of FXR. Although hepatic *Shp* expression was downregulated in VSG mice, additional FXR targets were either unchanged or upregulated. Furthermore, *Cyp7a1* mRNA levels were not consistently elevated across various cohorts of VSG mice, suggesting that transcriptional control of *Cyp7a1* may not play a role in promoting bile acid synthesis in our model. It also appears that ileal FXR-FGF15 signaling is not responsible for our results, as VSG mice exhibit increased bile acid synthesis in spite of comparable ileal *Fgf15* expression levels to sham mice. This is consistent with human studies, where serum C4 levels are elevated or unchanged post-VSG even in the presence of increased circulating FGF19 levels.^14,24^ These findings highlight an apparent disconnect or resistance to FXR and FGF15/19-mediated suppression of bile acid synthesis after VSG.

The lack of a significant weight difference between VSG and sham mice in our model is unexpected, since other murine models of VSG demonstrate sustained weight loss after surgery. One possible explanation includes our use of sutures to close the stomach after resection rather than surgical staples and clips, which may be somewhat restrictive. This is supported by the observation that VSG with staples leads to decreased food intake for 3-5 weeks after surgery and smaller, but more frequent meals.^25–27^ In contrast, VSG mice in our model only exhibited reduced intake between week 1-2, allowing VSG mice to catch up to their Sham counterparts at a faster rate. Additionally, it is unclear whether using the staple method may contribute to weight loss secondary to post-operative complications since mortality rates are not always reported. Our 2-week mortality rate was approximately 5% in sham vs 8.5% VSG mice. Any mouse found to have an abscess at the time of sacrifice was excluded from the study. Aside from the discrepancy in body weights and food intake, our study corroborates the observations of other groups. This includes intact absorption after VSG, as well as decreased hepatic steatosis despite similar body weights to sham.^18,28^ Notably, two other groups that use sutures have also observed that VSG mice do not display sustained weight loss but continue to retain the beneficial effects of the surgery.^29–30^

Our murine model of VSG lowers systemic and hepatic cholesterol levels independent of weight loss, providing a well-controlled and novel system to investigate how gastric resection influences hepatic physiology to reduce cholesterol burden. We have characterized the physiologic changes responsible for the decrease in cholesterol levels after VSG, but further studies are required to understand what may be driving bile acid synthesis after VSG and whether this process relies on CYP7A1 or alternative bile acid synthesis pathways. It is also unclear whether the effect of VSG on cholesterol metabolism contributes to an improvement in hepatic steatosis. Nonetheless, our findings are consistent with prior studies that have demonstrated that the beneficial effects of gastric resection go beyond food restriction and weight loss.

---

**Fig. S1.**
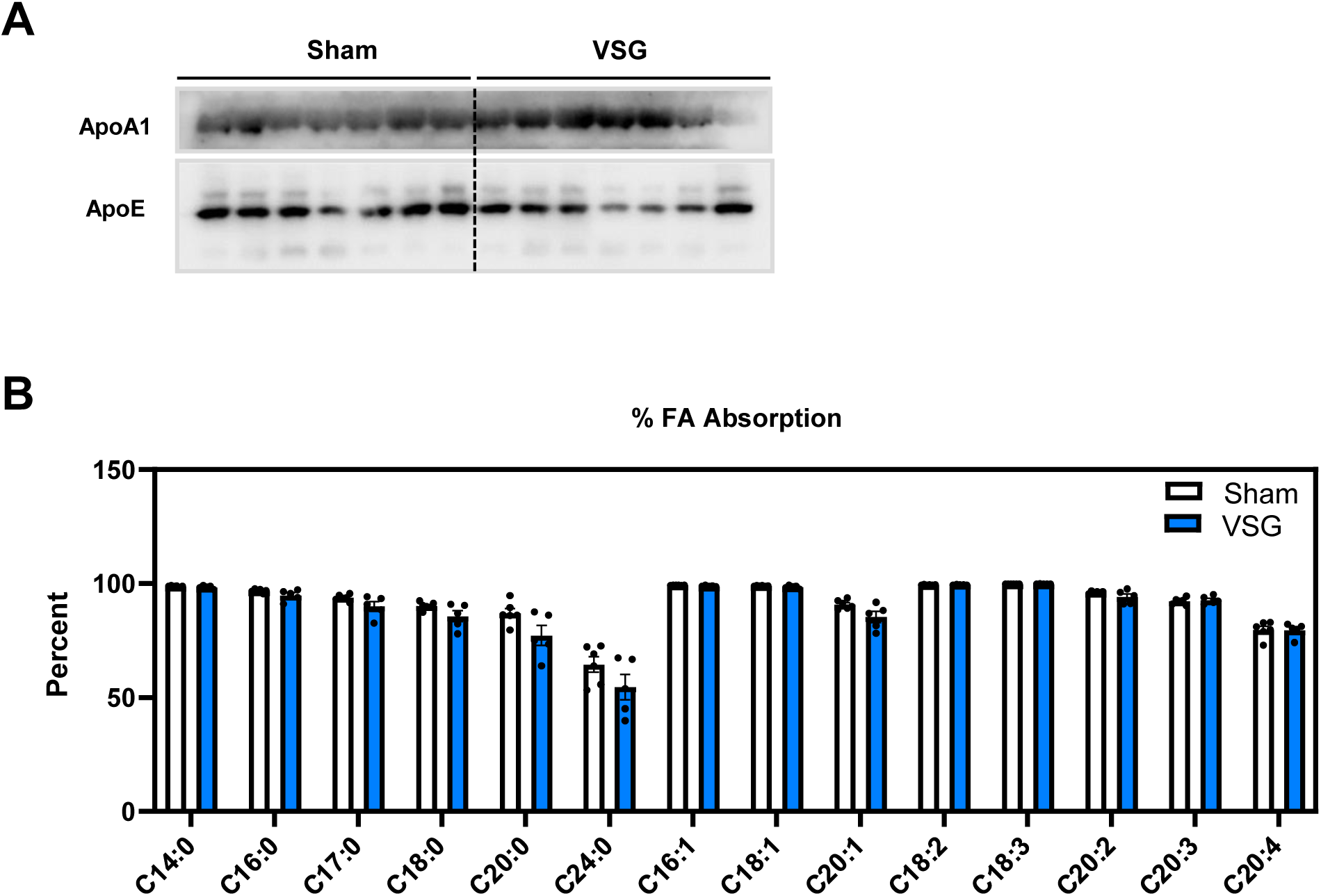
Fatty acid absorption is unaffected after VSG. (A) Week 6 fasting serum ApoA1 and ApoE in sham and VSG mice on WD; (B) Percent absorption of individual fatty acid species in sham and VSG mice on a 5% sucrose polybehenate diet at 6 weeks post-op, n = 6 vs 5.

**Fig. S2.**
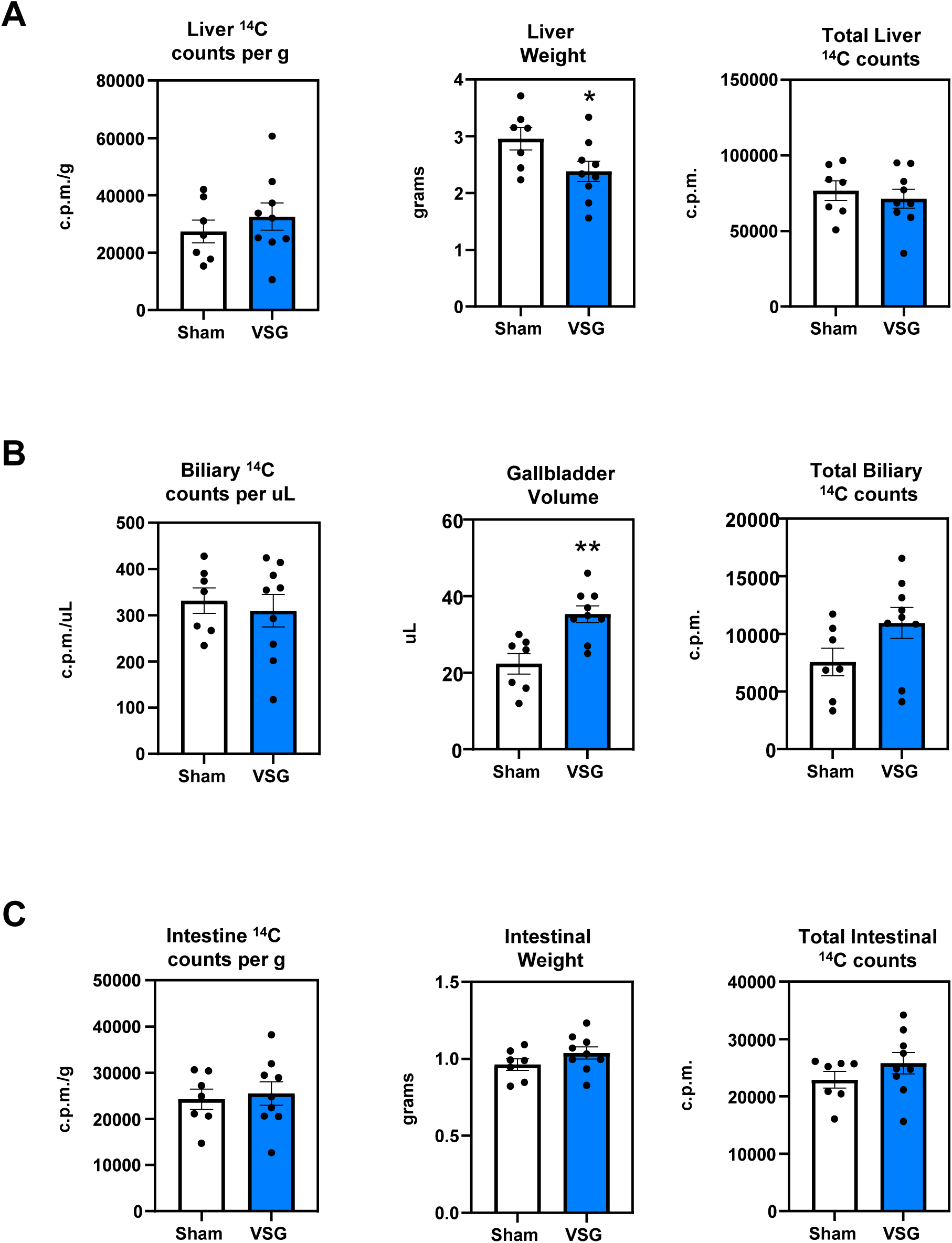
Distribution of [14C]-label 3 days after intravenous injection of [14C]-labeled HDL-cholesterol in the (A) liver, (B) gallbladder, and (C) intestine in sham and VSG mice at 7 weeks post-op, n = 7 vs 9. Expressed as 14C counts per gram or microliter and total 14C counts in the liver, gallbladder, and intestine. (*P < 0.05, ** P < 0.01, *** P < 0.001, ****P < 0.0001)

**Fig. S3.**
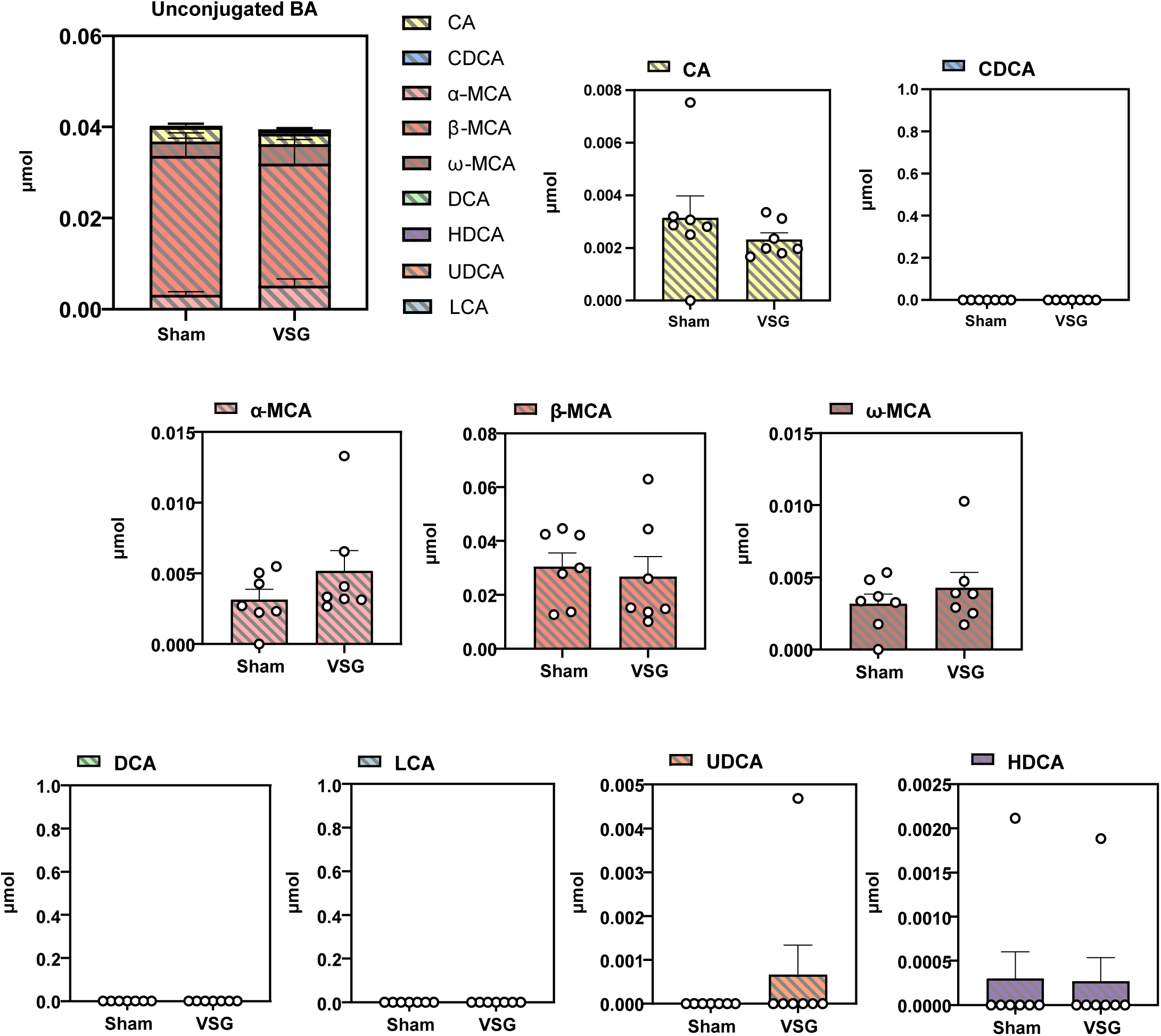
Unconjugated liver bile acids at 11 weeks post-op in sham and VSG mice, n = 7 vs 7. (*P < 0.05, ** P < 0.01, *** P < 0.001, ****P < 0.0001)

**Fig. S4.**
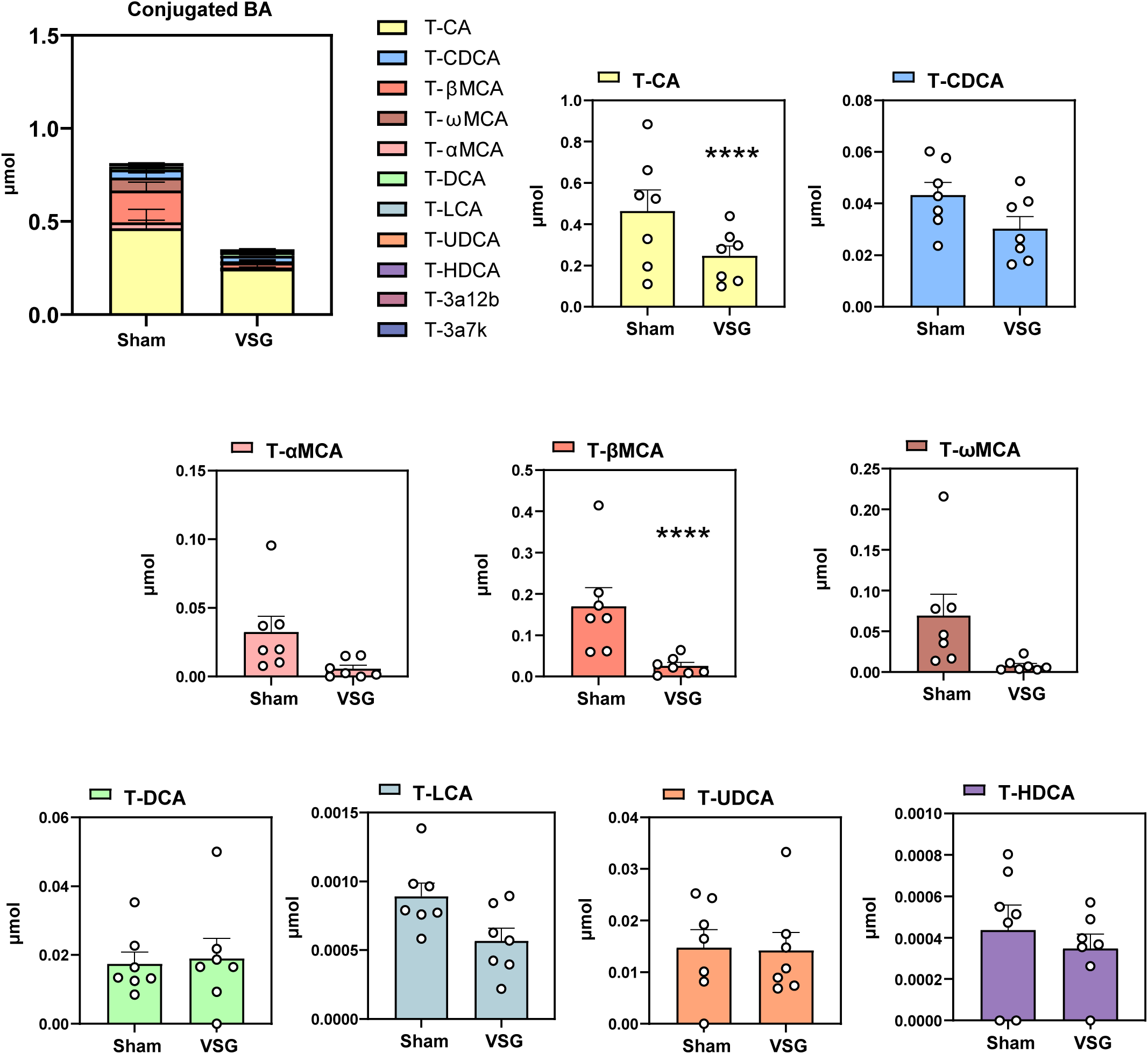
Conjugated liver bile acids at 11 weeks post-op in sham and VSG mice, n = 7 vs 7. (*P < 0.05, ** P < 0.01, *** P < 0.001, ****P < 0.0001)

**Fig. S5.**
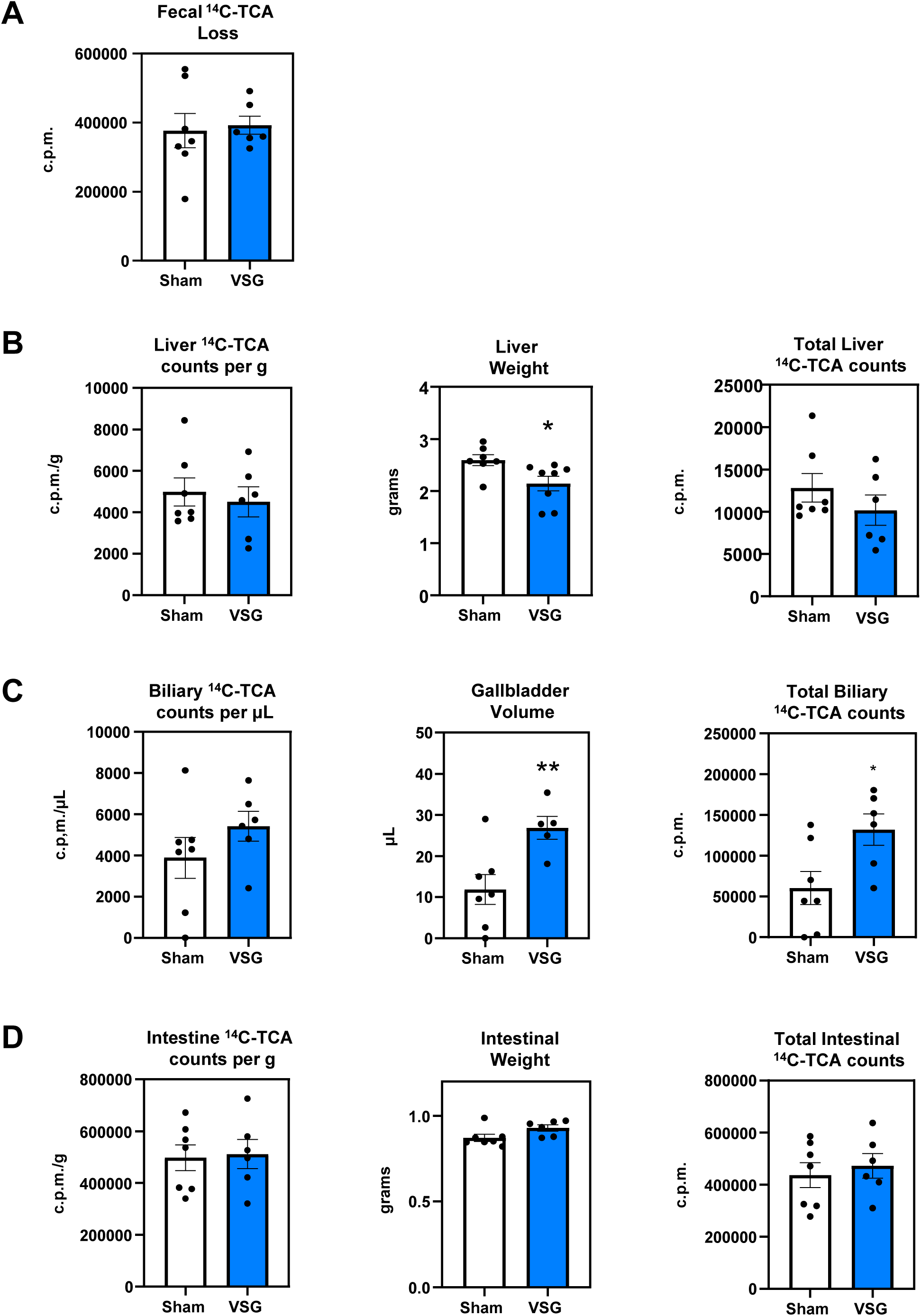
Metabolism of [14C]-labeled TCA 3 days after oral gavage of [14C]-labeled TCA at 10 weeks post-op. (A) Cumulative loss of 14C-TCA in feces over 3 days in sham and VSG mice, n = 7 vs 7. (B-D) Distribution of [14C]-TCA in the (B) liver, (C) gallbladder, and (D) intestine in sham and VSG mice, n = 7 vs 7. Expressed as 14C-TCA counts per gram or microliter and total 14C-TCA counts in the liver, gallbladder, and intestine. (*P < 0.05, ** P < 0.01, *** P < 0.001, ****P < 0.0001)

**Fig. S6.**
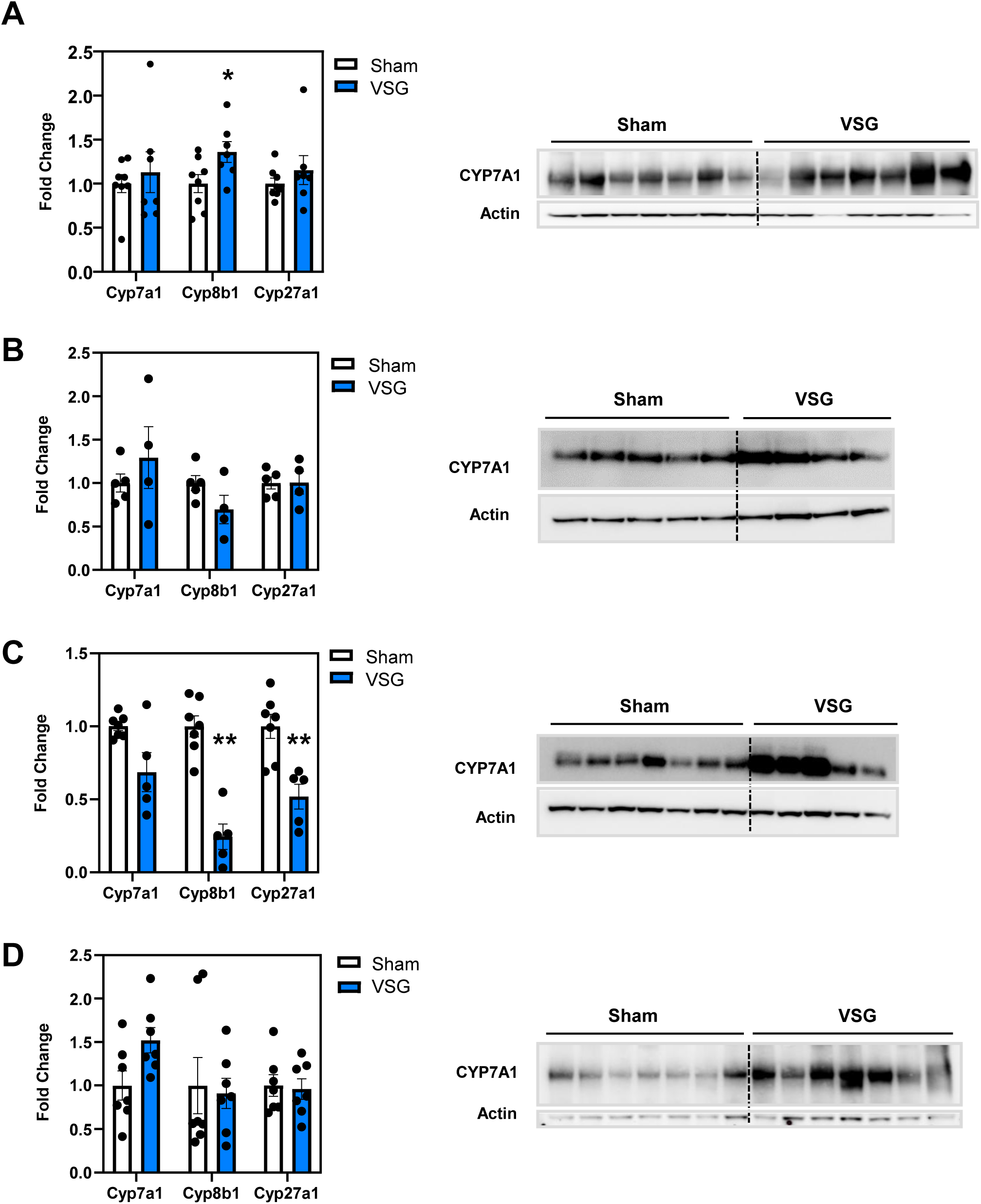
Hepatic Cyp7a1 gene expression changes with corresponding immunoblot in various cohorts of sham and VSG mice. (A) 11 weeks post-op on western diet after 16 hour fast; (B) 14 weeks post-op on western diet after 4 hour fast; (C) 10 weeks post-op on western diet after 4 hour fast; (D) 10 weeks post-op on western diet after 4 hour fast. (*P < 0.05, ** P < 0.01, *** P < 0.001, ****P < 0.0001)

## Notes

### Competing Interest Statement

The authors have declared no competing interest.

